# Integration of molecular coarse-grained model into geometric representation learning framework for protein-protein complex property prediction

**DOI:** 10.1101/2024.03.14.585015

**Authors:** Yang Yue, Shu Li, Yihua Cheng, Zexuan Zhu, Lie Wang, Tingjun Hou, Shan He

## Abstract

Structure-based machine learning algorithms have been utilized to predict the properties of protein-protein interaction (PPI) complexes, such as binding affinity, which is critical for understanding biological mechanisms and disease treatments. While most existing algorithms represent PPI complex graph structures at the atom-scale or residue-scale, these representations can be computationally expensive or may not sufficiently integrate finer chemical-plausible interaction details for improving predictions. Here, we introduce MCGLPPI, a novel geometric representation learning framework that combines graph neural networks (GNNs) with the MARTINI molecular coarse-grained (CG) model to predict overall PPI properties accurately and efficiently. This framework maps proteins onto a concise CG-scale complex graph, where nodes represent CG beads and edges encode chemically plausible interactions. The GNN-based encoder is tailored to extract high-quality representations from this graph, efficiently capturing the overall properties of the protein complex structure. Extensive experiments on three different downstream PPI property prediction tasks demonstrate that MCGLPPI achieves competitive performance compared with the counterparts at the atom- and residue-scale, but with only a third of the computational resource consumption. Furthermore, the CG-scale pre-training on protein domain-domain interaction structures enhances its predictive capabilities for PPI tasks. MCGLPPI offers an effective and efficient solution for PPI overall property predictions, serving as a promising tool for the large-scale analysis of biomolecular interactions.

## Introduction

Protein-protein interactions (PPIs) play a pivotal role in regulating diverse cellular processes, including signal transduction, immune response, and metabolic regulations^1,2^. Gaining insights into PPI aids in understanding protein functions and identifying potential drug targets^3–6^. While traditional experimental techniques for studying PPIs, such as yeast two-hybrid screening^7^, co-immunoprecipitation^8^, pull-down assays^9^, and fluorescence resonance energy transfer (FRET)^10^, are effective, they often require extensive labor and substantial financial investment. To address these challenges, advancements in computational tools and artificial intelligence (AI) algorithms have transformed the study of PPIs^11^. These in-silico strategies leverage expansive datasets to predict PPIs, enabling interaction site prediction^12^, interaction type classification^9^, and binding affinity prediction^13^.

The three-dimensional (3D) structures of proteins are fundamental to their biological functions^14–16^. To gain a nuanced understanding of the biological significance and detailed mechanisms underlying PPIs, decoding the geometry of protein complexes have become essential^1^. Among various computational methods, Graph neural networks (GNNs)^13,17^ stand out with their proficiency in handling the 3D structures of proteins. By integrating spatial information and topological data inherent to protein complexes, GNNs provide a robust framework for illuminating the multifaceted nature of protein interactions^18,19^. For instance, Jing et al.^20^ proposed a GNN framework, GVP-GNN, which preserves rotation equivariance of protein rigid motions when capturing geometric representations of protein-protein complexes. Zhang et al.^21^ designed a line-graph-augmented message passing scheme to inject the relative positional information between two interactive edges for different PPI prediction tasks, such as protein-protein interface identifications.

Notably, in GNNs-based methods, proteins are represented as graph structures, with nodes corresponding to either heavy atoms (i.e., the atom-scale model) or amino acids (i.e., the residue-scale model)^21,22^. However, each approach has its own trade-offs. Atom-scale models, while detailed, demand extensive computational resources to manage thousands of nodes, limiting their application to large PPI systems. On the other hand, residue-scale models are more computationally tractable but may overlook critical binding details that influence specificity and affinity. To address these limitations, multi-scale information can be integrated into the node features and edge connections. However, such integration require intricate information exchange across scales, maintaining model consistency and physical relevance, which can complicate the design process. Additionally, in both atom- and residue-scale models, edges typically represent interactions based on sequential threshold or geometric distance, aiming to capture the complex relationship between protein structures and functions. Nevertheless, using such criteria to define connections may misrepresent chemical bonds, potentially affecting predictive accuracy.

A potential solution to these issues is to adopt coarse-grained (CG) modeling, which is a well-established framework in protein molecular dynamics (MD) simulation, designed to concisely strike a balance between maintaining essential molecular details and enhancing computational efficiency. CG-scale representation simplifies groups of atoms into single sites, such as amino acid side chains or specific chemical groups. The MARTINI22 model^23^, a widely recognized CG-scale model in protein MD simulation, represents an average of four heavy atoms and their associated hydrogens with a single CG bead. It classifies beads into four main physical types, including polar (P), nonpolar (N), apolar (C), and charged (Q), with subtypes based on hydrogen bonding capabilities or polarity. In addition to the multiple bead types, the model includes numerous chemical-plausible interaction parameters, both bonded (bonds, angles and dihedrals) and nonbonded, to directly and accurately reflect the partitioning free energy of amino acid sidechains^24,25^. Through this strategy, the MARTINI model retains essential molecular interaction features while significantly reducing computational demands, making it invaluable for studying large protein complexes.

Although the CG-scale offers improved efficiency, its simulations still consume more resource than PPI predictions using AI techniques. Previous efforts to integrate the CG-scale model with machine learning (ML) or deep learning (DL) methods have primarily focused on optimizing force field potential parameters, predicting the peptide self-assembly shapes, and converting the CG-scale model back to atomistic structures^26,27^. However, a comprehensive approach that combines AI and CG modeling to predict PPI properties remains an under-explored area.

In this study, we presented MCGLPPI, a lightweight geometric representation learning framework that combines GNNs with the MARTINI22 CG-scale model to predict the overall properties of PPI complexes. Designed to optimize computational efficiency without compromising prediction accuracy, MCGLPPI employs a novel CG-scale complex graph, which maps each CG bead of protein complexes to nodes and utilizes chemical-plausible MARTINI force field bond parameters as edges for efficient structural characterization. Additionally, we introduced a GNN-based CG geometric-aware encoder to extract the high-quality representations from the devised graph.

Our extensive validation demonstrated that MCGLPPI achieved competitive performance on three of our curated overall property prediction benchmarks for PPIs, including two binding affinity prediction and one interaction type classification tasks. When compared to atom-scale and residue-scale counterparts, MCGLPPI significantly improved computational efficiency, and reduced graphics processing unit (GPU) usage and total running time by more than threefold without compromising accuracy. Moreover, proteins are intricate molecular machines that typically consist of multiple domains. Domain-domain interactions (DDIs) are critical subsets of PPIs, where the interaction typically occurs between domains rather than the entire proteins^28–30^. We demonstrated that the CG-scale pre-training based on DDI patterns significantly enhances the model’s ability to predict PPI binding affinity. Overall, MCGLPPI emerges as a general, accurate, and efficient method for predicting PPI properties, offering a pathway to sophisticated analysis of biomolecular interactions.

## Results

### Overall of MCGLPPI

We integrate biomolecular CG structures, force field parameters, and geometric-aware GNNs within the MCGLPPI framework for efficient prediction of overall properties of protein-protein complexes, which consists of three major components: (1) CG-scale complex graph generation, (2) CG-scale geometric representation learning, and (3) DDI-based CG-scale graph encoder pre-training. A comprehensive overview of the framework and its components is provided in Fig. 1.

**Fig. 1.**
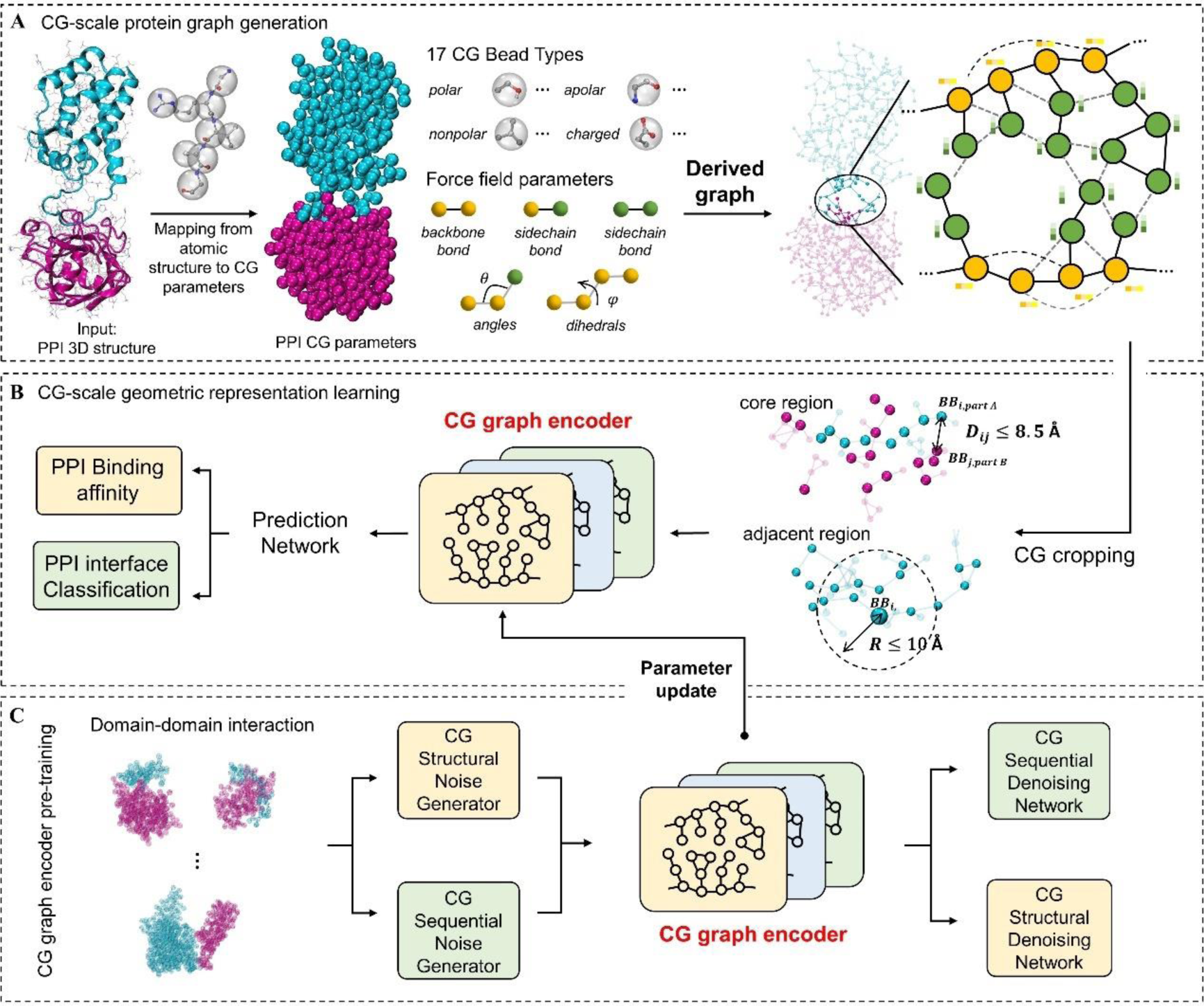
The flowchart of the MCGLPPI framework. **a,** CG-scale protein graph generation. The atomistic structure of a protein-protein complex is transformed into its corresponding coarse-grained (CG)-scale force field parameters using the MARTINI22 engine, with these parameters encompassing bead types and bonded interactions, which further include bonds, angles, and dihedrals. Based on these structural details and parameters, we create a CG-scale protein graph where beads are represented as nodes. Bonds between these beads are represented as edges in the graph, and information on angles and dihedrals is encoded as node features **b,** CG-scale geometric representation learning. The specifically designed CG-scale protein complex graph, containing comprehensive information of concise MARTINI beads and bonds, is firstly cropped to identify its core interaction region. The geometric representation of this cropped graph is extracted by the corresponding CG graph encoder, which is then fed into a protein prediction network to predict the overall property of the complex. **c,** DDI-based CG-scale graph encoder pre-training. The graph encoder can be better initialized by the CG-scale pre-training techniques applied to our carefully screened (domain-domain interaction) DDI dataset. After pre-training, the graph encoder with updated model parameters can be fine-tuned to generate geometric representations with potentially more powerful capability for downstream prediction tasks.

### Force field parameter and CG-scale complex graph generation

Structure-based prediction of PPI complex properties typically demands high-quality learning of protein geometric graph representations. The number of graph nodes and edges significantly affect computational cost. At the same time, it is crucial to ensure that the graph structure is chemically plausible, as it is essential for accurately depicting the properties of protein complexes.

On top of this, we introduce the CG scale-based MARTINI parameterization that aims to efficiently achieve a balanced representation between chemically plausible interaction characterization and computational cost. This process commences by transforming an atomistic PPI structure into a comprehensive set of CG-scale force field parameters tailored for the MARTINI22 model^23,31^. This simplification reduces the high-resolution atomic model into a computationally easier-to-execute form by grouping multiple atoms into fewer representative beads. The resulting concise interaction parameters are describe how these beads interact with each other chemically and physically from different perspectives (Fig. 2).

**Fig. 2.**
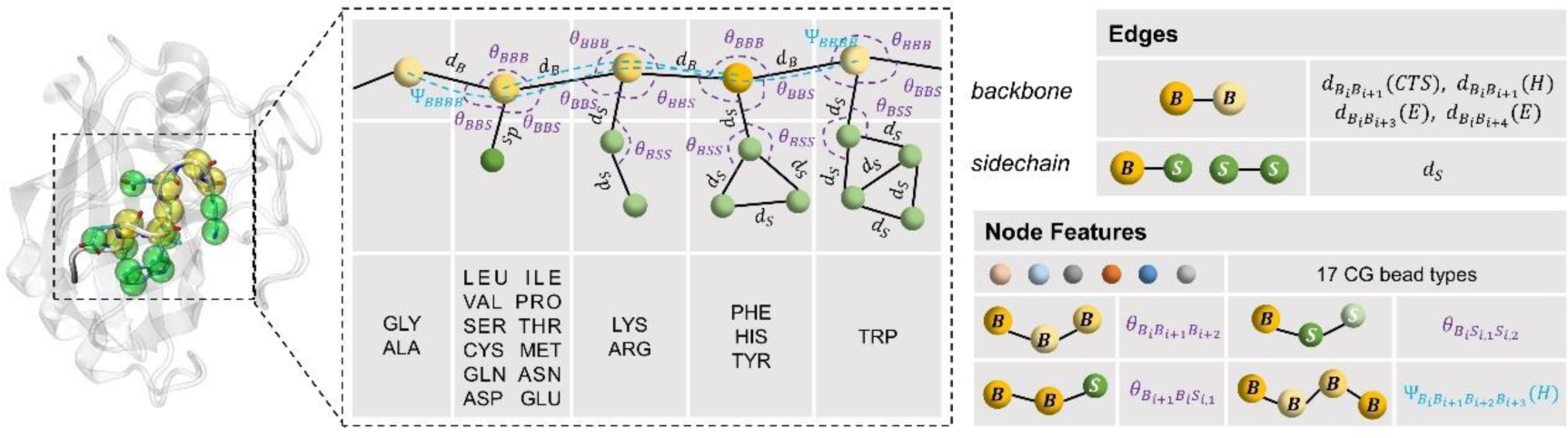
MARTINI22 CG-scale representation of protein structure. Each residue is represented by one backbone bead (*B*) and zero to four side-chain beads (*S*), depending on the residue type (left). The bonding interaction parameters within and between amino acids are shown (to effectively depict their local interaction structures), including bonds (types and lengths), angles (formed by triples of beads), and dihedrals (for quadruplets of beads). The right panel provides an in-depth view of the constructed CG-scale complex graph based on these concise parameters, showcasing both the edge connections and the node features.

After integrating the structural data with the force field parameters, a multi-relational graph corresponding to the protein complex is constructed (see Fig. 1a, Fig. 2, and Methods for further details). Within this graph, each bead, representing a group of heavy atoms, becomes a node. The bonds between backbone beads (*B*) or between sidechain (*S*) and either sidechain or backbone beads defined by their type and length, are translated into edges that connect these nodes. It is worth noting that these nodes and edges are concise (the corresponding statistics are provided in Supplementary information), ensuring efficient protein modeling while maintaining chemical accuracy. Within the MARTINI framework, the protein’s secondary structure plays a pivotal role in determining the bead types and associated bond, angle, and dihedral parameters for each residue. For instance, specific bond types such as constraint bonds *d*_*B*_*i*_*B*_*i+1*__ (*H*) or long harmonic bonds *d*_*B*_*i*_*B*_*i+3*__ (*E*) and *d*_*B*_*i*_*B*_*i+4*__ (*E*) are used for regions designated as helices (*H*) or extended strands (*E*), while other backbone bond parameters *d*_*B*_*i*_*B*_*i+1*__ (*CTS*) are adopted for irregular secondary structures such as coils, turns, and bends. In our CG-scale complex graph, the edge types also reflect these distinctions, facilitating the accurate description of secondary structural features within the protein complex. Furthermore, two distinct edge types, *d*_*intra*_ and *d*_*inter*_, detailed in the Methods section are introduced for the differentiation of bead nodes originating from the same or different amino acid residues, providing valuable hierarchical geometric information regarding the spatial arrangement relationships and interactions within and between the residues. Additionally, other crucial force field parameters, such as bead types, bond angles, and dihedrals, are encoded as node features within the graph (as illustrated in Fig. 2 and described in Methods). These features are essential for capturing the spatial orientation and potential movements of the protein segments.

Furthermore, as the MARTINI22 force field parameters of bond lengths, angles and dihedrals are derived from the statistical analysis of the Protein Data Bank (PDB)^24,32^ database, the construction of the CG-scale protein graph requires additional calibration and assignment of these parameters to accurately describe geometric interactive characteristics of protein complexes. Please refer to the Methods section for further details.

### CG-scale geometric representation learning

To reduce the computational overhead and preserve the integrity of the data for different PPI structures, we have implemented a duel-strategy approach to graph cropping on the CG-scale protein complex graphs derived earlier (Fig. 1b). The initial strategy, core region cropping, focuses on the interaction interface between two proteins (or defined interaction parts of the structures beyond dimers), by establishing a cut-off threshold for the backbone distance between any two amino acid residues. A residue is considered as part of the interaction interface if the distance between its backbone bead and that of a residue from the other protein/part is less than or equal to 8.5 Å. This criterion ensures that we focus on the most critical region of the interaction, likely enhancing model prediction accuracy and relevance. However, this core region cropping may omit peripheral but potentially significant structural information. To address this, we introduce an adjacent region cropping scheme, retaining any amino acid within a 10 Å backbone distance from the core region’s amino acids. This secondary measure captures essential spatially correlated motifs surrounding the core interface. By combining these strategies, we can produce a graph, that balances detailed structural information retention with computational feasibility, regardless of the interaction pattern. The specific cropping details are described in the Methods section.

We then apply the cropping method to each complex sample in our curated downstream datasets, which include two binding affinity regression and one interface type classification tasks. These tasks span a range of complexes, from the formation of simple dimers to the binding of the T cell receptor (TCR) to an antigenic peptide presented by the major histocompatibility complex (pMHC)^33,34^.

We subsequently utilize a multi-relational heterogeneous GNN-based CG graph encoder^21^, which can efficiently encode the complex relationships between graph nodes and edges (as detailed in the Methods section) to the cropped graph for generating its high-quality geometric representation. This representation is then forwarded to the task-specific prediction network, enabling us to obtain accurate predictions of the corresponding complex overall properties.

### DDI-based CG-scale graph encoder pre-training

Domains are fundamental structural units within proteins that are often responsible for specific functions. They play a critical role in mediating interactions with other proteins^28–30^, whether within a single multifaceted protein (intra-protein interactions) or between two distinct proteins (inter-protein interactions). Despite the limited availability of detailed yet labelled three-dimensional structural data for PPIs, the wealth of DDI structural information provides a valuable opportunity for enhancing computational models through pre-training. To this end, we use the Three-Dimensional Interacting Domains (3DID) database^28^ to construct a dataset tailored for pre-training our CG-scale graph encoder. The detailed curation process is described in the Methods section.

We employ a denoising-based, self-supervised pre-training approach, adapted from the work by Zhang et al.^22^, to instruct our CG graph encoder on the intricate patterns of DDI structures and sequences. This method involves introducing perturbation to each CG graph in the pre-training DDI dataset and then forcing the encoder to reconstruct the original graph information, thereby imprinting the fundamental characteristics of domain interactions (see Fig. 1c and the Methods section for more details). Following this pre-training phase, the encoder, now enriched with the knowledge from the DDI dataset, undergoes fine-tuning to tackle downstream PPI prediction tasks. Through this fine-tuning process, the encoder applies the principles of domain interactions learned during pre-training to downstream PPI scenarios, potentially enhancing its ability in making predictions.

### MCGLPPI saves computational cost while keeping competitive performance

To validate the performance and computational cost of the proposed MCGLPPI framework on the PPI complex overall property predictions, we curated three datasets: 1) the strict protein-protein dimer subset of the PDBbind dataset^35^ (PDBbind-strict-dimer dataset), 2) the ATLAS dataset^33^, and 3) the MANY/DC dataset^36,37^. The former two datasets were used to evaluate the model’s regression capabilities (protein-protein binding affinity predictions), while the MANY/DC dataset was used to assess the overall classification performance (protein-protein complex interface classifications). The Pearson’s correlation coefficient (*R*_P_), the root mean square error (RMSE), and the mean absolute error (MAE) were used to assess the quality of regressions. The area under the receiver operating characteristic curve (AUROC) and the area under the precision-recall curve (AUPR) were utilized the capability of classification. Additionally, to ensure a fair comparison, we performed the aforementioned complex graph cropping function for each sample in every dataset (across different scales) to identify the sample’s core interaction regions (detailed in Methods).

#### The binding affinity prediction of formation of strict dimers

We successfully extracted protein-protein complexes exhibiting strict dimer structures from the PDBbind dataset^35^. Following sample correction and label unification (specifically, converting the binding affinity labels of all relevant samples to Δ*G*)^38^, we obtained 1270 dimer samples with binding affinity labels Δ*G*, referred to as the PDBbind-strict-dimer dataset (the detailed curation process can be found in Supplementary Information). The tenfold cross-validation (CV) strategy was used to evaluate the model. To ensure a fair comparison of the model performance across different scales, we compared the atom- and residue-scale versions of our employed protein graph encoder, GearNet-Edge^21^, using their default model settings (i.e., settings related to protein graph construction and geometric encoder hyper-parameters). Besides, we considered an atom-scale state-of-the-art geometric encoder, GVP-GNN^20^, specifically designed for solving 3D macromolecular structures, particularly protein-protein complexes. Detailed information on the default hyper-parameters of all methods can be found in Supplementary Information.

Furthermore, to comprehensively quantify the cost of these approaches under limited lightweight computational resources, we utilized a single NVIDIA A100 GPU 40GB to run the comparative experiments. For each approach, based on the same epoch number of 150, starting with a batch size of 8 and gradually increased it by a factor of 2 until the GPU was out of memory (OOM), and we recorded the corresponding evaluation metrics, memory usage, and total time cost across the tenfold CV.

For the atom- and residue-scale GearNet-Edge, 915 of 1270 samples were successfully identified. To ensure a fair comparison, this 915-subset of the PDBbind-strict-dimer dataset was used for the comparative experiment. Table 1 presents the corresponding results. Under the current experimental conditions, the key findings were as follows: (1) MCGLPPI outperformed its atom- and residue-scale counterparts. (2) Under the same batch size, MCGLPPI reduced GPU consumption by approximately 5× and 3×, as well as total elapsed time by 3× and 3×, compared to the atom- and residue-scale models, respectively, while maintaining competitive performance. These findings demonstrated the effectiveness and feasibility of introducing the MARTINI-based mesoscopic CG-scale representation to achieve a better balance between performance and computational cost.

**Table 1.**
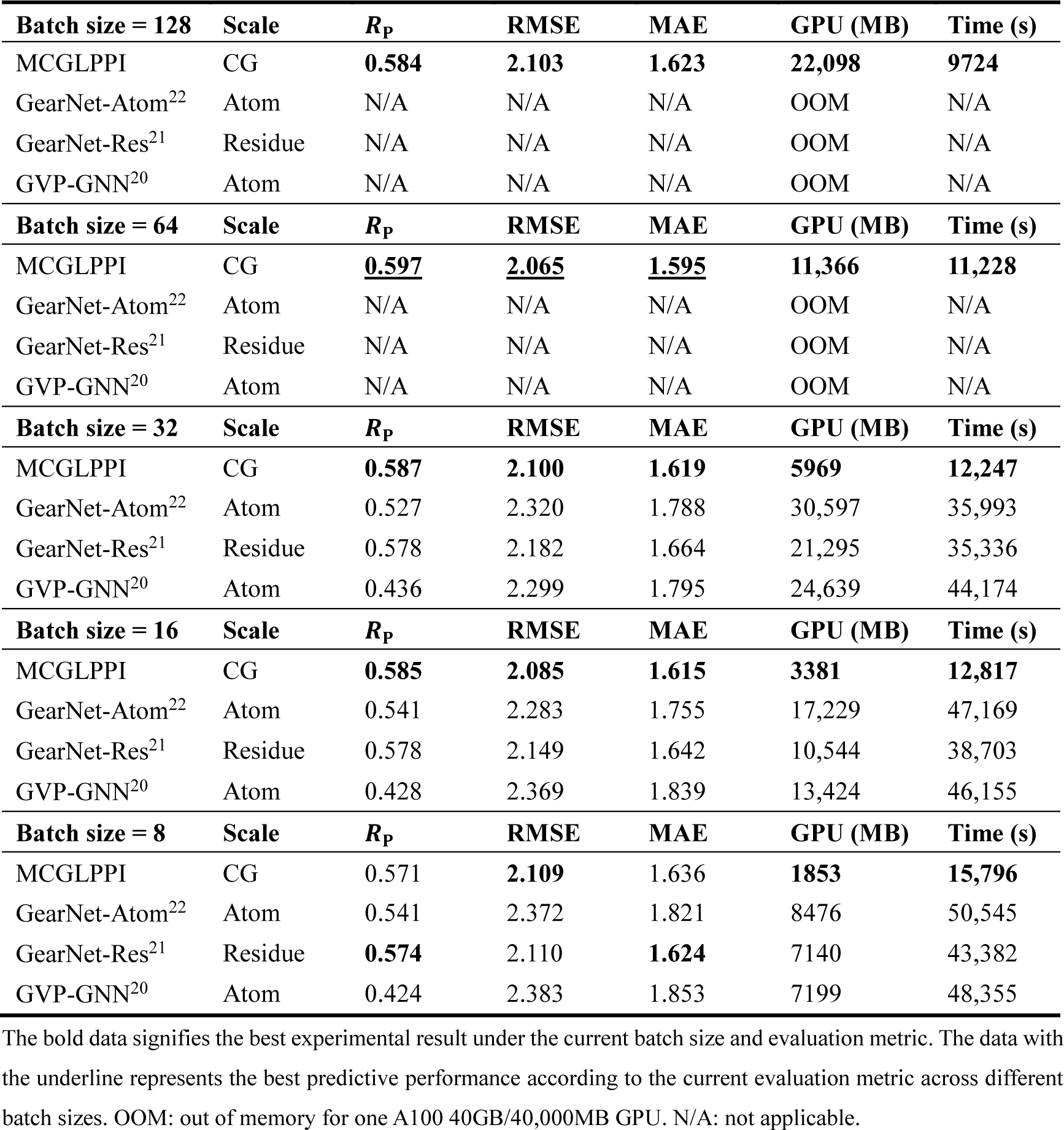
Test performance and computational cost of different approaches at different scales on the 915-subset of the PDBbind-strict-dimer dataset based on one A100 GPU 40GB. The default model settings of atom-scale GearNet-Edge (here denoted as GearNet-Atom), residue-scale GearNet-Edge (denoted as GearNet-Res), and (atom-scale) GVP-GNN were adopted from refs. ^22, 21^, and ^20^, respectively.

We also presented the experimental results of MCGLPPI on the complete PDBbind-strict-dimer dataset: 0.590 (*R*_P_), 2.071 (RMSE), 1.602 (MAE), 11,560 (GPU (MB)), and 14,312 (Time (s)). Besides, to validate the robustness of our proposed MCGLPPI, we conducted further investigations into the impact of hyper-parameter, such as hidden feature dimensions, on overall performance (see Supplementary Information). These investigations consistently supported the conclusions outlined above.

#### The effectiveness of MCGLPPI on more complex protein-protein interaction patterns

To further investigate the effectiveness of MCGLPPI in handling more complex protein-protein interaction structures beyond standard dimers, the ATLAS dataset^33^, which contains the TCR-pMHC structures formed in the cell-mediated immunity processes along with their corresponding binding affinity values, was considered. After removing invalid samples, correcting samples, and unifying labels, we obtained 531 different structures with the Δ*G* labels. Please note that we utilized the structures that were optimized using the fixed backbone design option of Rosetta^39^, which were reported to achieve high structural accuracy^33^.

We performed the tenfold cross-validation using the same experimental settings as the previous section and documented the corresponding evaluation results. Furthermore, the comprehensive comparison experiments were carried out based on 451 of the 531 curated samples that could be effectively processed by GearNet-Edge at both the atom- and residue-scale.

Table 2 shows the predictive performance and computational cost derived from the tenfold cross-validation performed on the 451-sample ATLAS subset. Additionally, we reported the best-performing results of MCGLPPI on the complete curated ATLAS dataset: 0.809 (*R*_P_), 1.116 (RMSE), 0.837 (MAE), 13,615 (GPU (MB)), and 6982 (Time (s)). Notably, when dealing with more complex protein-protein structures beyond standard dimers, the proposed MCGLPPI maintained the competitive performance and exhibited a relatively lower computational cost compared with its atom- and residue-scale counterparts, which further validated the effectiveness of the devised CG-scale protein complex geometric model and corresponding cropping function. An additional investigation into the necessity of the cropping function is conducted in the section of

**Table 2.** Test performance and computational cost of different approaches at different scales on the 451-subset of the curated ATLAS dataset based on one A100 GPU 40GB. The default model settings of atom-scale GearNet-Edge (here denoted as GearNet-Atom), residue-scale GearNet-Edge (denoted as GearNet-Res), and (atom-scale) GVP-GNN were adopted from refs. ^22, 21^, and ^20^, respectively.

| Batch size = 128 | Scale | $R_p$ | RMSE | MAE | GPU (MB) | Time (s) |
| --- | --- | --- | --- | --- | --- | --- |
| MCGLPPI | CG | **0.807** | **1.084** | **0.833** | **26,358** | **5578** |
| GearNet-Atom<sup>22</sup> | Atom | N/A | N/A | N/A | OOM | N/A |
| GearNet-Res<sup>21</sup> | Residue | N/A | N/A | N/A | OOM | N/A |
| GVP-GNN<sup>20</sup> | Atom | N/A | N/A | N/A | OOM | N/A |
| Batch size = 64 | Scale | $R_p$ | RMSE | MAE | GPU (MB) | Time (s) |
| MCGLPPI | CG | **0.825** | **1.054** | **0.782** | **13,285** | **5915** |
| GearNet-Atom<sup>22</sup> | Atom | N/A | N/A | N/A | OOM | N/A |
| GearNet-Res<sup>21</sup> | Residue | N/A | N/A | N/A | OOM | N/A |
| GVP-GNN<sup>20</sup> | Atom | N/A | N/A | N/A | OOM | N/A |
| Batch size = 32 | Scale | $R_p$ | RMSE | MAE | GPU (MB) | Time (s) |
| MCGLPPI | CG | **0.820** | **1.061** | **0.778** | **6796** | **5992** |
| GearNet-Atom<sup>22</sup> | Atom | 0.813 | 1.109 | 0.851 | 31,018 | 21,375 |
| GearNet-Res<sup>21</sup> | Residue | 0.818 | 1.068 | 0.796 | 21,059 | 19,866 |
| GVP-GNN<sup>20</sup> | Atom | 0.524 | 1.599 | 1.169 | 26,232 | 27,231 |
| Batch size = 16 | Scale | $R_p$ | RMSE | MAE | GPU (MB) | Time (s) |
| MCGLPPI | CG | **0.821** | **1.042** | 0.788 | **3524** | **7225** |
| GearNet-Atom<sup>22</sup> | Atom | 0.816 | 1.062 | **0.772** | 16,155 | 22,579 |
| GearNet-Res<sup>21</sup> | Residue | 0.820 | 1.056 | 0.816 | 11,314 | 22,276 |
| GVP-GNN<sup>20</sup> | Atom | 0.534 | 1.494 | 1.111 | 13,211 | 30,027 |
| Batch size = 8 | Scale | $R_p$ | RMSE | MAE | GPU (MB) | Time (s) |
| MCGLPPI | CG | 0.818 | **1.048** | **0.801** | **1892** | **8790** |
| GearNet-Atom<sup>22</sup> | Atom | **0.823** | 1.106 | 0.813 | 8255 | 26,810 |
| GearNet-Res<sup>21</sup> | Residue | 0.809 | 1.086 | 0.832 | 6190 | 24,157 |
| GVP-GNN<sup>20</sup> | Atom | 0.480 | 1.573 | 1.170 | 6946 | 32,253 |

#### The prediction results for protein-protein interface classification

In addition to the aforementioned two regression tasks, an overall interface classification task for protein-protein complex was incorporated to further examine the generalizability of MCGLPPI. Specifically, the MANY^36^ and DC^37^ datasets were utilized, containing 5739 and 161 dimers respectively. These dimers are categorized into two overall types: dimers with biological or crystal interfaces^40^. Based on this classification, the model was trained to distinguish between the two interface types, which was further formulated as a binary (complex graph) classification task. Following the previous data splitting convention^1,18^, 80% of MANY, 20% of MANY, and the complete DC datasets were used as the training, validation, and test sets for model evaluation, respectively.

The experiment settings from the previous two sections were maintained (except for the unified epoch number changing from 150 to 30). Additionally, we compared our approach with two existing approaches, DeepRank-GNN^18^ and EGGNet^1^, which had already been tested on the complete MANY/DC dataset. However, it should be noted that the effective sample numbers for atom- and residue-scale GearNet-Edge on the MANY and DC datasets were 5535 and 151, respectively. Moreover, the node feature construction in existing approaches like DeepRank-GNN relies on time-consuming external amino acid (AA) sequence alignment search, making it difficult to fairly compare computational cost. Therefore, we only compared their predictive performance on the complete MANY/DC dataset and conducted detailed computational cost comparison experiments for the atom- and residue-scale GearNet-Edge models on the 5535-151-sample subset (following the aforementioned data splitting mode).

The results of the computational cost comparison experiments are shown in Table 3. It was observed that, compared to its atom- and residue-scale counterparts, MCGLPPI achieved lower computational cost while surpassed their predictive capability. Specifically, MCGLPPI (with batch size 32) outperformed atom-scale GearNet-Edge by 3.9% (AUROC: 0.883 vs. 0.850) and residue-scale GearNet-Edge by 1.5% (AUROC: 0.883 vs. 0.870). Similarly, MCGLPPI outperformed atom-scale GearNet-Edge by 6.3% (AUPR: 0.876 vs. 0.824) and residue-scale GearNet-Edge by 2.0% (AUPR: 0.876 vs. 0.859). The reason for this improvement is attributed to the integration of protein thermodynamics and specific secondary structure support information through the MARTINI force field, which is injected into the bonds (edges) of the CG complex graph, which provides extra distinguishable capability compared with its atom- and residue-scale counterparts. A further investigation into the importance of different CG graph edges are presented in the **Performance of the geometries considered in CG-scale complex graphs** section.

**Table 3.** Test performance and computational cost of different approaches at different scales on the 5535-151-subset of the MANY/DC dataset based on one A100 GPU 40GB. The default model settings of atom-scale GearNet-Edge (here denoted as GearNet-Atom), residue-scale GearNet-Edge (denoted as GearNet-Res), and (atom-scale) GVP-GNN were adopted from refs. ^22, 21^, and ^20^, respectively.

| Batch size = 128 | Scale | AUROC | AUPR | GPU (MB) | Time (s) |
| --- | --- | --- | --- | --- | --- |
| MCGLPPI | CG | **0.875** | <u>**0.880**</u> | **25,437** | **584** |
| GearNet-Atom<sup>22</sup> | Atom | N/A | N/A | OOM | N/A |
| GearNet-Res<sup>21</sup> | Residue | N/A | N/A | OOM | N/A |
| GVP-GNN<sup>20</sup> | Atom | N/A | N/A | OOM | N/A |
| Batch size = 64 | Scale | AUROC | AUPR | GPU (MB) | Time (s) |
| MCGLPPI | CG | <u>**0.890**</u> | **0.871** | **13,269** | **617** |
| GearNet-Atom<sup>22</sup> | Atom | N/A | N/A | OOM | N/A |
| GearNet-Res<sup>21</sup> | Residue | N/A | N/A | OOM | N/A |
| GVP-GNN<sup>20</sup> | Atom | N/A | N/A | OOM | N/A |
| Batch size = 32 | Scale | AUROC | AUPR | GPU (MB) | Time (s) |
| MCGLPPI | CG | **0.883** | **0.876** | **6737** | **702** |
| GearNet-Atom<sup>22</sup> | Atom | 0.842 | 0.824 | 34,452 | 3055 |
| GearNet-Res<sup>21</sup> | Residue | 0.848 | 0.842 | 24,368 | 3031 |
| GVP-GNN<sup>20</sup> | Atom | 0.850 | 0.845 | 27,467 | 4132 |
| Batch size = 16 | Scale | AUROC | AUPR | GPU (MB) | Time (s) |
| MCGLPPI | CG | **0.860** | **0.849** | **3952** | **951** |
| GearNet-Atom<sup>22</sup> | Atom | 0.850 | 0.799 | 19,290 | 3293 |
| GearNet-Res<sup>21</sup> | Residue | 0.844 | 0.843 | 13,505 | 3208 |
| GVP-GNN<sup>20</sup> | Atom | 0.817 | 0.808 | 15,506 | 4515 |
| Batch size = 8 | Scale | AUROC | AUPR | GPU (MB) | Time (s) |
| MCGLPPI | CG | **0.881** | 0.857 | **2229** | **961** |
| GearNet-Atom<sup>22</sup> | Atom | 0.839 | 0.758 | 10,026 | 4017 |
| GearNet-Res<sup>21</sup> | Residue | 0.870 | **0.859** | 8578 | 3661 |
| GVP-GNN<sup>20</sup> | Atom | 0.836 | 0.825 | 8427 | 5194 |

Furthermore, we observed that the performance of MCGLPPI (AUROC: 0.895, AUPR: 0.892) also outperformed both DeepRank-GNN (AUROC: 0.865, AUPR: 0.871) and EGGNet (AUROC: 0.869, AUPR: 0.863) on the complete MANY/DC dataset (the results of DeepRank-GNN and EGGNet were retrieved from the refs ^1,18^). This observation further supported the effectiveness and generalizability of the MCGLPPI framework for predicting overall properties of protein-protein interaction complexes.

### The investigation of CG-scale pre-training techniques on different tasks

To explore the feasibility of our hypothesis that pre-training on CG-scale informative domain-domain interaction (DDI) complexes could benefit the downstream property predictions of CG complexes, especially in scenarios with limited labeled samples, we constructed a large dataset from the 3DID database^28^, which comprises 41,663 representative DDI structures, serving as our pre-training repository. These DDI structural samples were then transformed into CG-scale graphs. Furthermore, we implemented a CG-scale diffusion denoising-based self-supervised pre-training technique based on ref ^22^, which enables us to capture and learn the general DDI patterns and knowledge.

Specifically, we first determined the optimal model settings for MCGLPPI under each downstream task through training from scratch separately. Using the same selected settings (for each task), we fine-tuned the CG graph encoder that had undergone pre-training for each respective downstream task (with the same epoch numbers as training from scratch). We then compared the performance difference between training from scratch and pre-training with fine-tuning. As shown in Fig. 3a, for the PPI binding affinity prediction tasks on the two datasets, PDBbind and ATLAS, pre-training improved the *R*_P_ from 0.597 to 0.610 and from 0.825 to 0.836, respectively, indicating that pre-training can effectively enhance model performance. However, we also observed that for the interface type classification task on the MANY/DC dataset, the performance actually decreased after pre-training, with AUPR dropping from 0.880 to 0.865 (additional results evaluated using other metrics were reported in Supplementary Information).

**Fig. 3.**
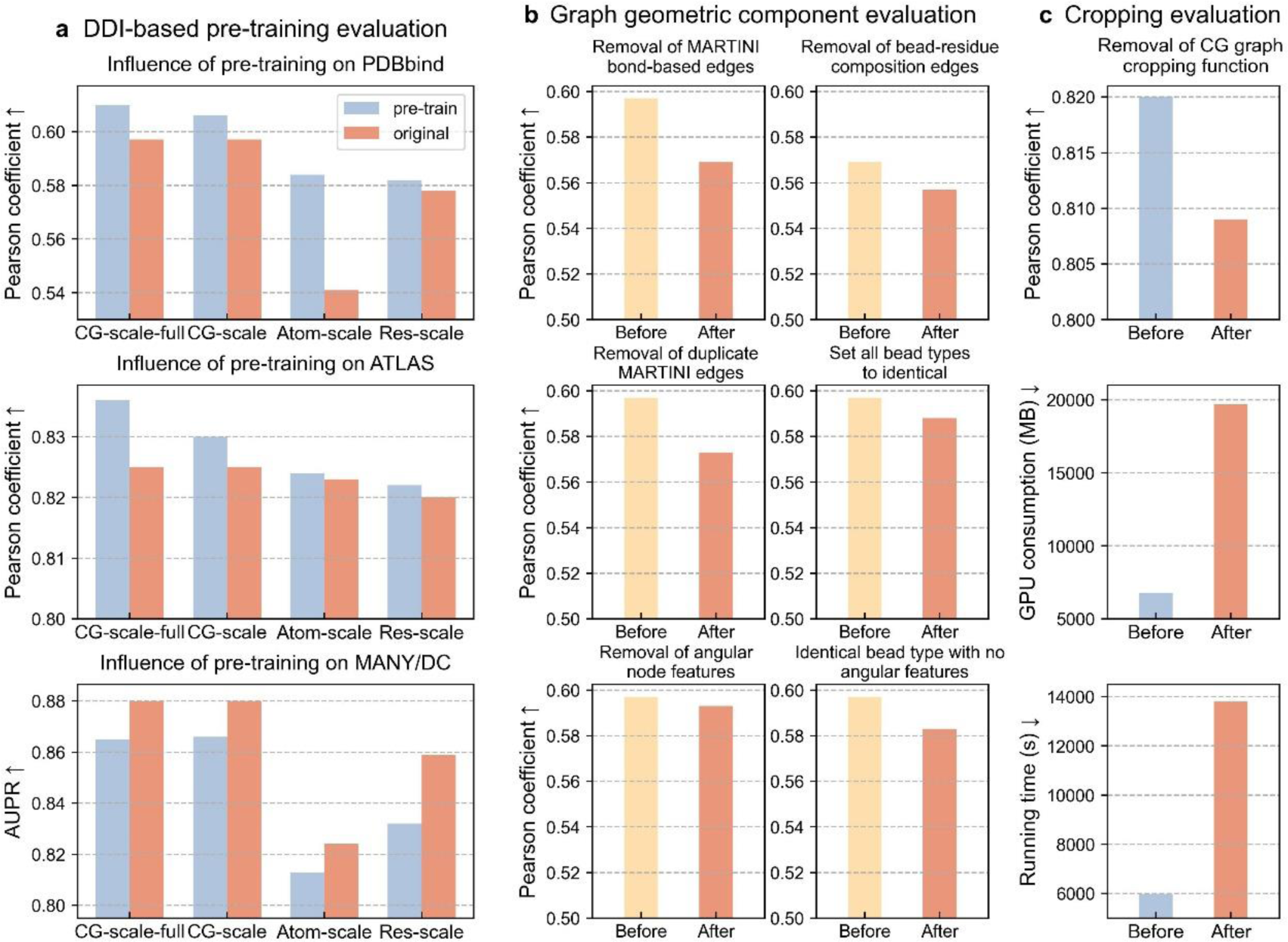
The investigation of performance influence contributed by the DDI-based CG-scale pre-training, crucial graph geometric characterization components, and cropping function of MCGLPPI. **a,** Performance influence brought by the DDI diffusion denoising-based pre-training on the three downstream datasets. “CG-scale-full”, “CG-scale”, “Atom-scale”, “Res-scale” represent the CG-scale pre-training on the complete 3DID pre-training set, CG-scale pre-training on the 33144-subset, atom-scale pre-training on the 33144-subset, and residue-scale pre-training on the 33144-subset, respectively; and the blue and red bars display the results obtained after and before imposing the pre-training, respectively. **b,** Performance influence of removing different geometric characterization components from the devised CG-scale complex graph. The respective experiments about the removal of different components (based on PDBbind) are indicated on top of each sub-table, and the yellow and red bars give the results acquired before and after removing the corresponding components, respectively. **c,** Performance influence related to the complex cropping function. Based on the maximum reachable batch size of MCGLPPI without the complex cropping function (i.e., 32), the performance comparison results of MCGLPPI before (blue bars) and after (red bars) closing the cropping function based on the ATLAS dataset, including the *R*_P_, GPU consumption, and total tenfold CV running time are provided.

We thought the contributing reasons are as follows. For the binary classification task that aims to distinguish between biological interfaces and crystal artefacts (representing non-biological interactions), training from scratch with current graph settings might be sufficient for the protein learning model to capture the subtle geometric structural difference between these interface types. Furthermore, pre-training based on DDI complexes extracted from actual (biological) protein-protein interactions would be more beneficial for tasks which predict the properties of complexes formed through real PPI processes (e.g., binding affinity predictions), rather than distinguishing crystallographic interfaces resulting from non-biological interactions detected between repetitive crystal units^18^.

To further explore the potential of the DDI dataset-based pre-training in enhancing PPI prediction models, we extended our performance comparison to the atom- and residue-scale counterpart encoders (i.e., pre-training them using the original-scale corresponding pre-training settings^22^ based on the 3DID pre-training set). It should be noted that the atom- and residue-scale models (i.e., GearNet-Edge) could only recognize 33,144 out of the 41,663 DDI samples. Therefore, we selected these 33,144 samples as an additional pre-training subset and transformed these samples into their respective atom-, residue-, and CG-scale graphs to compare the performance across different scales before and after pre-training.

Consistent with the results of the CG graph encoder pre-trained on the full set of DDI samples, the models at all three scales showed improved performance with 33144-sample-based pre-training compared to those without pre-training in the PDBbind and ATLAS datasets. However, in the MANY/DC dataset for crystallographic interface classifications, the model’s performance at all three scales also deteriorated after the pre-training. Additionally, when using the same number of pre-training samples, our CG-scale approach exhibited the overall best performance on the PDBbind and ATLAS datasets. In total, pre-trainings on DDIs are effective in enhancing PPI predictions, with the CG-scale graph encoder combined with CG DDIs pre-training being the most effective approach.

### Performance of the geometries considered in CG-scale complex graphs

To assess the impact of various geometric representations within CG complex graphs, we conducted a series of ablation studies. These experiments were designed to gradually remove specific graph components and analyze their individual contributions to the overall predictive performance of the system.

Initially, we focused on edges based on chemically plausible interactions as defined by the MARTINI force field. These interactions included various types of bonds between beads, such as *d*_*B*_*i*_*B*_*i+1*__ (*CTS*), *d*_*B*_*i*_*B*_*i+1*__ (*H*), *d*_*B*_*i*_*B*_*i+3*__ (*E*), *d*_*B*_*i*_*B*_*i+4*__ (*E*), and *d*_*S*_ (detailed in Methods). We used the subset of 915 protein dimers from the PDBbind-strict-dimer dataset for our analysis. Upon selective removal of all MARTINI bond-based edges from the CG graphs, we observed a measurable decline in performance metrics. The *R*_P_ decreased from 0.597 to 0.569 (Fig. 3b), confirming the importance of these edges in accurately characterizing protein interactions.

Next, after removing MARTINI bond-based edges, we investigated the effectiveness of the proposed bead-residue geometric hierarchical composition-aware edges *d*_*intra*_ and *d*_*inter*_ (detailed in Methods). We replaced these two types of edges with the standard radius-based edges that do not differentiate the compositional relationships between chemically plausible bead nodes and their corresponding residues (using the same cutoff). This modification resulted in a further decrease in predictive accuracy, with *R*_P_ dropping from 0.569 to 0.557 (Fig. 3b). This emphasized the significant impact that composition-aware edges have on the model.

We also evaluated scenarios within a complete CG graph where, if both residue composition-aware edges and MARTINI bond-based edges are present between the same pair of end nodes, the MARTINI bond-based edges are disregarded. Under these conditions, the predictive performance experienced a decline from an *R*_P_ of 0.597 to 0.573 (Fig. 3b). This result emphasized the importance of explicitly incorporating MARTINI bond-based edges along with the composition-aware edges in our modeling framework.

Furthermore, other MARTINI force field parameters, such as bead types, angles (*θ*_*B*_*i*_*B*_*i+1*_*B*_*i+2*__, *θ*_*B*_*i+1*_*B*_*i*_*S*_*i,1*__, *θ*_*B*_*i*_*S*_*i,1*_*S*_*i,2*__), and dihedrals (*Ψ*_*B*_*i*_*B*_*i*+1_*B*_*i+2*_*B*_*i+3*__), were encoded as node features within the graph. To assess their importance, we invalidated the bead type feature and angular features, which include (bond) angles and dihedrals, from the CG-scale graph. As shown in Fig. 3b, when bead types for each node were set to identical or when angular information was omitted, there was a reduction in the *R*_P_ from 0.597 to 0.588 and 0.593, respectively. Moreover, the complete invalidation of node features related to the MARTINI force field, including both bead type and angular features, led to a further decline in *R*_P_ to 0.583. These findings indicated that, not only does our framework rely on aforementioned chemical-plausible edges, but it also requires both the chemical and physical information provided by bead types and the angular information from angles and dihedrals. This combined information enables MCGLPPI to make more informed predictions about the properties of protein interactions.

### Influence of graph cropping on overall model efficiency

To provide a comprehensive analysis of the influence of graph cropping on the efficiency of the MCGLPPI framework, we conducted additional experiments using the ATLAS dataset, which contains more complex TCR-pMHC structures as a benchmark. We ran our model on the subset of 451 protein complex structures from the ATLAS dataset while maintaining all experimental settings consistent with our previous MCGLPPI experiments, except for the graph cropping function, which was disabled. As shown in Fig. 3c, the maximum batch size that could be processed by MCGLPPI under a single NVIDIA A100 GPU 40G decreased significantly from 128 to 32 when the graph cropping function was turned off. With a batch size of 32, the *R*_P_, GPU memory consumption, and total runtime post-cropping were 0.825, 6796 MB, and 5992 seconds, respectively, compared to 0.809, 19,706 MB, and 13,794 seconds before cropping. These results confirmed the critical role of graph cropping in improving computational efficiency and predictive performance in our framework.

## Discussion

In this study, we presented MCGLPPI, a novel framework that enhances the structure-based overall property predictions for protein-protein complexes by utilizing the MARTINI force field for lightweight protein modeling. At the mesoscopic CG-scale, our proposed CG protein graph model uses concise yet chemically plausible beads and bonds to accurately represent the conformation characteristics of protein-protein complexes. This approach results in lower computational overhead, leading to a better balance between predictive performance and cost compared to atom- and residue-scale models.

Furthermore, our extensive ablation studies highlighted the significance of both edge and node features derived from the MARTINI force field for accurate PPI prediction using the MCGLPPI framework. Notably, these features, particularly chemical bond-based edges and physical-plausible bead type node features, are crucial in capturing the essential properties that govern protein interactions. Additionally, through our proposed CG-scale learning framework, we demonstrated the effectiveness of DDI-based pre-training in improving binding affinity predictions of PPIs.

While MCGLPPI has shown promising overall performance, there are some areas for further improvement or investigation. For instance, MCGLPPI may not fully capture the complexity of protein-protein systems, and other mesoscopic CG-scale protein modeling systems deserve further exploration. We plan to incorporate more cost-effective geometric information to more comprehensively characterize CG complex structures, e.g., considering the Euler angles to describe the relative rotation relationships between CG particles, and integrate more CG modeling systems to capture protein-protein thermodynamic quantities and underlying chemical mechanisms from different perspectives.

## Methods

### The detailed curation process for the 3DID pre-training dataset

The latest 3DID database^28^ provides 15,983 DDI structure templates, with each template containing one or more samples of resolved 3D structural data. To prevent data leakage, we removed any DDI templates from the 3DID dataset that were identical to those present in our downstream benchmark datasets. Following this stringent exclusion process, we obtained a robust pre-training dataset which provides 41,663 DDI structure samples in total.

### MARTINI-based geometric parameter generation

Each complex structure was processed using pdbfixer tool (https://github.com/openmm/pdbfixer) to complete missing side-chain information and convert non-natural amino acids to their natural counterparts (for the atom- and residue-scale models, the same process was also performed). We utilized the martinize.py Python script to convert an atomistic PPI structure into a set of MARTINI22 CG-scale force field parameters, which can be further categorized into two types of parameters, i.e., structure and topology parameters. The former ones encompass the bead coordinate-related information, while the latter ones further include both nonbonded parameters, such as bead types, and bonded parameters, including bonds, angles, dihedrals, and bead connectivity instructing the bead composition for these bonds and angles (Fig. 2).

#### MARTINI-based bonds information

Within the MARTINI22 force field representation, there are two principal types of bonds: backbone bonds and sidechain bonds. As illustrated in Figs. 1a and 2, a backbone bond *d*_*B*_ is formed between two neighboring backbone beads (*B*_*i*_). Sidechain bonds *d*_*S*_ occur either between a backbone bead and a sidechain bead (*S*_*i*_) within the same amino acid or between sidechain beads. Furthermore, the MARTINI force field differentiates backbone bond types based on the protein’s secondary structure. Consequently, *d*_*B*_ can be subdivided into:

1. Constraint bonds: These are formed between two adjacent amino acids that are part of a helical structure (*H*), denoted as *d*_*B*_*i*_*B*_*i+1*__ (*H*).
2. Long harmonic backbone bonds: For three consecutive amino acids forming extended elements (*E*), these bonds connect the backbone beads of residues *i* and *i* + 3, denoted as *d*_*B*_*i*_*B*_*i+3*__ (*E*).
3. Long harmonic backbone bonds: For four contiguous amino acids in extended elements (*E*), the bonds connect backbone beads of residues *i* and *i* + 4, denoted as *d*_*B*_*i*_*B*_*i+4*__ (*E*).
4. Other harmonic backbone bonds: Parameters between two adjacent amino acids for irregular secondary structures such as coils, turns, and bends are denoted as *d*_*B*_*i*_*B*_*i+1*__ (*CTS*).

In total, there are five types of bonds in the MARTINI22 force field: *d*_*B*_*i*_*B*_*i+1*__ (*CTS*), *d*_*B*_*i*_*B*_*i+1*__ (*H*), *d*_*B*_*i*_*B*_*i+3*__ (*E*), *d*_*B*_*i*_*B*_*i+4*__ (*E*), and *d*_*S*_. These bond types were chosen as the primary edge types in our MCGLPPI framework.

#### MARTINI-based angles and dihedrals information

The bonded parameters also include angle and dihedral parameters (Fig. 2). Specifically, there are three types of angle parameters and one dihedral type being considered:

1. *θ*_*B*_*i*_*B*_*i+1*_*B*_*i+2*__ : The angle between three consecutive backbone beads.
2. *θ*_*B*_*i*+1_*B*_*i*_*S*_*i,1*__ : The angle formed between a backbone bead, its neighboring backbone bead, and the first sidechain bead of the amino acid.
3. *θ*_*B*_*i*_*S*_*i,1*_*S*_*i,2*__ : The angle between a backbone bead and two consecutive sidechain beads of the same amino acid.
4. *Ψ*_*B*_*i*_*B*_*i*+1_*B*_*i+2*_*B*_*i+3*__ : The dihedral angle between four consecutive backbone beads. It is noted that dihedral angles were imposed only when all four interacting beads had the helical secondary structure (*H*) in MARTINI force field.

Notably, the bond lengths, angle and dihedral values were re-calibrated based on the given coordinates of corresponding endpoint beads. They were not directly adopted from the statistics values from PDB database since the calibration can provide accurate geometric interactive information for the CG-scale protein complex graph model.

### The construction of CG-scale protein complex graph and its cropping function

The challenge to build an effective CG protein complex graph is how to fully preserve the introduced MARTINI parameters in a graph structure accurately and efficiently, while keeping the flexibility of injecting other useful knowledge on top of these MARTINI parameters. To overcome this challenge, we first modelled a given protein-protein complex as a multi-relational contact graph *G* = (*V, E, R*). *V* represents the set of graph nodes *i*, i.e., all MARTINI beads produced for the complex, and the position of each bead node *i* is determined by its equipped 3D coordinate. ℰ and ℛ are the set of edges between bead nodes and the set of edge types *r*, respectively. Based on this, we denoted an edge from nodes *j* to *i* with type *r* as (*i*, *j*, *r*)).

In order to give the precise descriptions of inter-bead geometric positional relationships while integrating concise chemical-plausible MARTINI bonds, the graph structure is built as follows. First, an edge will be wired if any two bead nodes have the Euclidean distance smaller than 5Å. Compared with the commonly-used radius edge used in atom- and residue-scale models, MCGLPPI further distinguishes the edge type based on whether the two end bead nodes are from the same residue. In other words, the edge is categorized as an intra-residue contact edge *d*_*intra*_ if the two bead nodes belong to the same residue otherwise is an inter-residue contact edge *d*_*inter*_, for which the hierarchical composition information between chemical-plausible beads and their surrounding residues within a complex is injected. Besides, all bond types described in **MARTINI-based bonds information** are incorporated in the graph edge structure. To summarize, there are seven types of edges in total, describing the protein complex geometry from different perspectives, including various precise geometric contact relationships, actual secondary structure supports, and chemical bonded interactions, etc. Although multiple types of edges are included, due to the conciseness of MARTINI bonds, the average degree of nodes in the constructed graph is still relatively small (see Supplementary Information for the further comparison analysis with the atom- and residue-scale counterparts), which contributes to relatively low representation learning overhead. Next, the other MARTINI parameters were allocated as follows into these defined nodes and edges as their features.

*Bead node features f*_*i*_.

- The MARTINI22 bead type, given as a one-hot representation
- Sine-cosine encoded backbone angles ([*sin*(*θ*_*BBB*_), *cos*(*θ*_*BBB*_)])
- Sine-cosine encoded backbone-side chain angles ([*sin*(*θ*_*BBS*_), *cos*(*θ*_*BBS*_)])
- Sine-cosine encoded side chain angles ([*sin*(*θ*_*BSS*_), *cos*(*θ*_*BSS*_)])
- Sine-cosine encoded backbone dihedrals ([*sin*(*Ψ*_*BBBB*_), *cos*(*Ψ*_*BBBB*_)])

*Graph edge features f*_(*i*,*j*,*r*)_.

- The one-hot MARTINI22 bead type of the source node
- The one-hot MARTINI22 bead type of the target node
- The edge type, given as a one-hot representation
- The absolute positional difference between source and target nodes in the MARTINI22 bead sequence, given as a one-hot representation
- The calibrated bond length

For the above individual node and edge features, they were concatenated as the final features (*f*_*i*_: *R* ∈ 1 × 25, *f*_(*i*,*j*,*r*)_: *R* ∈ 1 × 53). Besides, since the angular parameters generated from MARTINI are sparse (see Fig. 2, not every bead will involve in the calculation of every type of angles), an extra rule detailed in Supplementary Information was provided, to assign sparse angular node features to specific beads to avoid potential conflicts.

After the definition of the CG-scale complex graph, the corresponding graph cropping function was designed to identify its core interaction regions for further reducing the computational cost and potentially increasing the predictive accuracy. Specifically, we first determined the interaction parts of the protein-protein complexes with different interaction patterns. For the curated complexes in the PDBbind and MANY/DC datasets, they belong to the standard dimers, we chose each protein chain as one part, thus a complex can be represented as two interaction parts. For the structures in ATLAS which usually contain 4 or 5 chains, the peptide and MHC chains were treated as the first part, while the second part contained the remained TCR chain structures.

Next, a distance matrix *M*^*dis*^ based on the specified two interaction parts was created to guide the generation of the cropped complex graph *G*^′^. For each residue, MARTINI will only assign one backbone bead (*B*) with a 3D coordinate to represent its backbone atoms and their overall position, and thus the coordinate of *B* was used as the position of the residue (analogous to using alpha carbon (*Cα*) as the residue position in residue-level protein graph constructions^21^). Based on this, *M*^*dis*^ with the size of *L*_*AA*1_ × *L*_*AA*2_ can be calculated, in which *L*_*AA*1_ and *L*_*AA*2_ are the AA sequence length (given by above backbone beads *B*) of the interaction parts 1 and 2, respectively. In *M*^*dis*^, every element 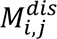 was the pairwise Euclidean distance between corresponding residues from individual interaction parts (given by *B*_*i*_ and *B*_*j*_). Then any pair of residues having the distance smaller than 8.5Å were retained as the core region. Other AAs that had *B* − *B* distance to any core region AAs smaller than 10Å were also retained.

After that, all bead nodes and edges within the retained region were kept as the cropped complex graph *G*^′^ (the angular node features were further re-calibrated if any end bead nodes within the angular information calculations were removed by this cropping). Furthermore, for the atom-scale and residue-scale models, to conduct a fair comparison, the similar cropping function were employed to their protein graphs, the only difference was replacing *B* with *Cα* to indicate the overall position of the residue.

### The CG-scale representation learning for complex overall property prediction

After acquiring the cropped protein complex graph *G*^′^, a representative multi-relational heterogeneous GNN-based protein encoder GearNet-Edge^21^, was incorporated into the framework, for predicting the overall property of protein complexes. Specifically, based on a line graph-enhanced edge message passing mechanism^41^ to model the inter-edge positional relationships, the additional structural information can be injected into the node representations for more effective protein geometric interaction modeling (the corresponding equations are provided in Supplementary Information). We made it work at the CG-scale for generating the overall geometric representation of the input CG cropped graph *G*^′^, and the generated representation was further learnt by a three-layer task-specific multi-layer perception (MLP) to give the final property prediction result.

### The DDI-based CG graph encoder pre-training technique

The protein domain-domain complex parameterized by MARTINI still preserves the fundamental conformation and chain sequence, but the basic particles are substituted from original atoms to CG beads. Intuitively, performing the self-supervised noise-adding-denoising pre-training techniques, which are already demonstrated to be effective on understanding geometric regulations of proteins at atom- or residue-scale^42^, could also benefit the understanding of general knowledge from CG DDI complexes (for downstream property predictions).

Therefore, based on the atom-scale work^22^, a CG-scale complex pre-training technique was developed, which adds noise with changing magnitudes into 3D coordinates and sequences of MARTINI-based CG bead nodes based on the diffusion mechanisms^43^, for the CG-complex geometric regulation learning. The equations and complete details are provided in Supplementary Information. After the pre-training, the trained CG graph encoder will be fine-tuned on the specified downstream task to produce the effective structural representations for corresponding input CG complex graphs.

For the implementation of the CG-scale protein complex graph construction, representation learning, and pre-training processes, Pytorch^44^ and Torchdrug^45^ with a default random seed 0 was employed, and the Adam^46^ with the initial learning rate of 0.0001 was adopted as the optimizer for model training (the same environment settings were also used for the other involved models working at the atom- and residue-scale). Besides, all experiments were deployed on a configuration of one NVIDIA A100 GPU 40GB. An complete summary of tools for implementing MCGLPPI is given in Supplementary Information.

## Code availability

The source code of MCGLPPI can be downloaded from https://github.com/arantir123/MCGLPPI.

## Data availability

The input source data of MCGLPPI can be found in https://github.com/arantir123/MCGLPPI. All other data that supports the results of this study is available from the corresponding author upon request.

## Acknowledgements

This work was supported by the Computer Science Ramsay Fund at the University of Birmingham. We are thankful for the financial support from Macao Polytechnic University Foundation (RP/FCA 07/2022 to S.L.). We also thank Mr. D. McDonald for his helpful suggestions on our task studies.

## Author contributions

Y.Y. and S.L. conceived the research project. Y.C. and S.H. supervised the research project. Y.Y. designed the computational pipeline. Y.Y. implemented the MCGLPPI framework and performed the model training and prediction validation tasks. S.L. curated all involved protein complex samples and conducted experiments for MARTINI force field-based geometric parameter generation. Y.Y., S.L., Y.C., T.H., and S.H. wrote the manuscript with support from all authors.

