## Supplementary Material for "Integration of molecular coarse-grained model into geometric representation learning framework for protein-protein complex property prediction"

### The statistics of graph nodes and edges of the MANY/DC dataset

Based on the standard dimer dataset MANY/DC[1,2] and conventional dataset splitting settings[3,4], we give the statistics of nodes and edges of the protein complex graphs at different scales, in which the atom- and residue-scale complex graphs are constructed based on the default settings of atom- and residue-scale GearNet-Edge, respectively[5,6] (as the counterparts of the MCGLPPI).

Supplementary Table 1: the statistics of nodes and edges in each set of the MANY/DC dataset

| **Scale** | **Sub-set** | **Graph node number** | **Graph edge number** | **Average Node degree** |
|---|---|---|---|---|
| CG | Training | 321.780 | 2373.738 | 7.377 |
| CG | Validation | 321.315 | 2371.637 | 7.381 |
| CG | Test | 392.377 | 2900.821 | 7.393 |
| **Scale** | **Sub-set** | **Graph node number** | **Graph edge number** | **Average Node degree** |
| Atom | Training | 1226.182 | 18,897.831 | 15.412 |
| Atom | Validation | 1221.720 | 18,807.455 | 15.394 |
| Atom | Test | 1504.132 | 23,402.179 | 15.559 |
| **Scale** | **Sub-set** | **Graph node number** | **Graph edge number** | **Average Node degree** |
| Residue | Training | 156.996 | 3023.984 | 19.261 |
| Residue | Validation | 156.553 | 3016.193 | 19.266 |
| Residue | Test | 192.682 | 3772.550 | 19.579 |

We can conclude that, compared with the atom-scale counterpart based on the full-atom characterizations, the CG-scale protein complex graph produced by MCGLPPI has overall smaller graph size. Compared with the residue-scale counterpart which edges are fully built based on the pre-defined geometric distance and sequential thresholds, the edges in the CG graph are more concise and chemical-plausible, which ultimately contribute to overall more sparse and accurate edge distribution for efficient inter-particle interaction descriptions.

### Detailed curation process of the PDBbind-strict-dimer dataset

We retrieve 2852 protein–protein complexes with known binding affinity data in total from the PDBbind database (version 2020). Initially, we refine this dataset to include only the simplest types of complexes, which are composed of two protein components.

We then select those samples that form a single PPI binding interface, as determined by their three-dimensional (3D) structural configurations. In cases where a two-component complex has multiple PPI binding interfaces, but these interfaces are structurally similar, we retain the sample and extract the structural information for the two proteins forming one representative binding interface. These filtering procedures result in a strict dataset of 1270 dimeric complexes (termed as the PDBbind-strict-dimer dataset).

### The default hyper-parameters of involved approaches in comparison

We compare our CG-scale MCGLPPI framework with the counterpart GearNet-Edge which can function at the atom- or residue- scales[5,6]. We also incorporate an atom-scale advanced approach GVP-GNN[7], which is specifically designed to learn 3D macromolecular structures (esp., protein-protein complexes), into the complete comparison experiments. We give the basic default hyper-parameters of these involved approaches from original papers[5-7] as follows.

1. MCGLPPI (CG-scale): The detailed graph construction settings are provided in Methods of the manuscript. The default CG-scale graph convolutional layer number and hidden feature dimension are 6 and 256, respectively.
2. GearNet-Edge (atom-scale)[5,6]: The construction of protein graph structures for GearNet-Edge at the atom-scale is based on the radius edge: each atom node $i$ will connect all other atom nodes within a 4.5Å-radius 3-dimensional (3D) sphere centered at $i$. The default graph convolutional layer number and hidden feature dimension are 6 and 128, respectively.

3. GearNet-Edge (residue-scale)[5,6]: The construction of protein graph structures for GearNet-Edge at the residue-scale is built based on three different types of distance-based or sequence-based edges.
    1) Radius edge: Analogous to those used in GearNet-Edge (atom-scale), the only difference is that the radius cutoff is set to 10 Å when wiring the neighboring residue nodes.
    2) KNN edge: Each residue node $i$ connects other 10 nodes nearest to it, based on the Euclidean distance between their node position coordinates (and an extra 5 Å cutoff is imposed for limiting the minimum inter-residue distance).
    3) Sequential edge: Two residue nodes will be wired if these two nodes have the relative positional difference $N$ in the amino acid (AA) sequence of current protein. The pre-defined $N$ includes 1, 2, and 3.

    Besides, The default graph convolutional layer number and hidden feature dimension are 6 and 512, respectively.
4. GVP-GNN (atom-scale)[7]: Based on the same radius edge setting as GearNet-Edge at the atom-scale (4.5Å cutoff), the graph structure is constructed. GVP-GNN explicitly distinguishes scalars and vectors within the node and edge features. For the default hidden dimensions for node features, 100 (for scalars) and 16 (for vectors) are adopted. For the hidden dimensions of edge features, 32 (for scalars) and 1 (for vectors) are chosen. Besides, its default graph convolutional layer number is set to 5.

### Investigation about the influence of hidden feature dimensions on

### overall performance of MCGLPPI

Because the default hidden feature dimensions of MCGLPPI and its atom- and residue-scale counterparts (i.e., GearNet-Edge) are 256, 128, and 512, respectively (the default graph convolutional layer number is the same: 6). In order to investigate the robustness of our MCGLPPI to the default dimensions of its atom- and residue-scale counterparts, we conduct an extra experiment as follows.

Specifically, based on the experimental results on 915-subset of the PDBbind-strict-dimer dataset (see Table 1 of the manuscript), we can find the top predictive performance of atom- and residue-scale GearNet-Edge is both achieved at batch size equaling to 16. Therefore, we separately adjust the hidden feature dimension of MCGLPPI into 128 and 512 at the batch size 16 to conduct a fair comparison, and the results under the corresponding settings are recorded.

For the combination of hidden feature dimension 128 plus batch size 16, the results are: 0.587 ($R_{\mathrm{P}}$), 2.088 (RMSE), 1.615 (MAE), 1625 (GPU (MB)), and 9818 (Total Time (s)). The corresponding results under the combination of hidden feature dimension 512 plus batch size 16 are: 0.579 ($R_{\mathrm{P}}$), 2.079 (RMSE), 1.614 (MAE), 6658 (GPU (MB)), and 24,893 (Total Time (s)). We can observe that, after keeping the same hidden feature dimensions as the atom- and residue- counterparts, MCGLPPI still achieves the competitive predictive performance while preserving relatively lower computational cost (refer Table 1 of the manuscript, and the optimal results of MCGLPPI are achieved at a combination of hidden feature dimension 256 with batch size 64). This can validate the robustness of MCGLPPI to the hidden feature dimension, and further demonstrate the effectiveness of our CG-scale protein geometric representation learning framework.

### The experimental results based on other metrics for the evaluation of DDI-based pre-training technique

Following the same experimental settings described in **The investigation of CG-scale pre-training techniques on different tasks** section of the manuscript, we here provide the predictive performance of the pre-trained MCGLPPI and its atom- and residue-scale counterparts, which is evaluated based on the other metrics (except for $R_P$ and AUPR) on the downstream datasets.

From the experimental results (Table S2), we find that under the current pre-training settings and evaluation metrics, the MCGLPPI still outperforms its atom- and residue-scale counterparts, which is in line with the results shown in Fig. 3a of the manuscript.

Supplementary Table 2: the other experimental results of the pre-training at different scales

| **PDBbind** | | | | |
|---|---|---|---|---|
| **Pre-training set** | **Model name** | **Scale** | **RMSE** | **MAE** |
| Complete 3did | MCGLPPI | CG | **2.037** | **1.572** |
| The 33144-subset | MCGLPPI | CG | 2.053 | 1.588 |
| The 33144-subset | GearNet-Edge | Atom | 2.163 | 1.646 |
| The 33144-subset | GearNet-Edge | Residue | 2.165 | 1.632 |
| **ATLAS** | | | | |
| **Pre-training set** | **Model name** | **Scale** | **RMSE** | **MAE** |
| Complete 3did | MCGLPPI | CG | **0.998** | 0.765 |
| The 33144-subset | MCGLPPI | CG | 1.002 | **0.745** |
| The 33144-subset | GearNet-Edge | Atom | 1.052 | 0.771 |
| The 33144-subset | GearNet-Edge | Residue | 1.056 | 0.808 |
| **MANY/DC** | | | | |
| **Pre-training set** | **Model name** | **Scale** | **AUROC** | |
| Complete 3did | MCGLPPI | CG | 0.874 | |
| The 33144-subset | MCGLPPI | CG | **0.877** | |
| The 33144-subset | GearNet-Edge | Atom | 0.838 | |
| The 33144-subset | GearNet-Edge | Residue | 0.855 | |

The bold data indicates the best experimental result under current dataset and evaluation metric.

### Detailed description of allocation of sparse angular features to specific bead nodes

For each angular parameter generated by the MARTINI22 engine, its sine-cosine encoded feature will be allocated to specific bead node based on the following rules:

- The backbone angles ($\theta_{BBB}$): For each $\theta_{BBB}$, the corresponding encoded feature will be assigned to the second bead node of current $BBB$ combination.
- The backbone-side chain angles ($\theta_{BBS}$): For each $\theta_{BBS}$, the corresponding encoded feature will be assigned to the third bead node of current $BBS$ combination.
- The side chain angles ($\theta_{BSS}$): For each $\theta_{BSS}$, the corresponding encoded feature will be assigned to the third bead node of current $BSS$ combination.
- The backbone dihedrals ($\Psi_{BBBB}$): For each $\Psi_{BBBB}$, the corresponding encoded feature will be assigned to the second bead node of current $BBBB$ combination.

### The equations of the GearNet-Edge protein encoder

Concisely, GearNet-Edge introduces a line graph-augmented edge message passing strategy[8], which models the inter-edge relative positional relationship, to inject the additional structural information into the node representations for more effective protein geometric characteristics learning. The relevant equations are as follows:

$$h_i^0 = f_i \tag{1}$$

$$h_i^l = h_i^{l-1} + u_i^l \tag{2}$$

$$u_i^l = \sigma(BN\left(\sum_{r\in\mathcal{R}} W_r \sum_{j\in N_r(i)} \left(h_j^{l-1} + MLP\left(m_{(i,j,r)}^l\right)\right)\right)) \tag{3}$$

$$m_{(j,i,r_1)}^0 = f_{(j,i,r_1)} \tag{4}$$

$$m_{(j,i,r_1)}^l = \sigma(BN(\sum_{r\in\mathcal{R}'} W_r' \sum_{(k,w,r_2)\in N_r'((j,i,r_1))} m_{(k,w,r_2)}^{l-1})) \tag{5}$$

in which $f_i$ and $f_{(j,i,r)}$ are initial node and edge features, $h_i^l$ and $u_i^l$ are the hidden representation of node $i$ at the $l$-th (graph convolutional) layer and aggregated neighboring information of node $i$ at the $l$-th layer, respectively. For the calculation of $u_i^l$ (equation (3)), $\sigma$, $BN$, $W_r$, $N_r(i)$, $MLP$, and $m_{(i,j,r)}^l$ represent the Sigmoid activation function, Batch Normalization layer[9], edge-type-specific linear transformation matrix, 1-hop neighbors of node $i$ with edge type $r$, multi-layer perception (MLP), and augmented edge feature based on the line graph at $l$-th layer, respectively. Furthermore, for the calculation of the augmented edge feature (equation (5)), it follows the similar information aggregation mechanism as equation (3), which will aggregate the neighboring edges' information into the feature of the central edge based on a pre-defined line graph. For the description of the line graph construction among existing edges, please refer Zhang et al.[5] for detailed information.

### The details about the CG-scale pre-training technique

For every protein domain-domain (DDI) complex curated from the 3DID dataset, we first transform it into a CG-scale complex graph following the same procedure as that for downstream complex transformations. These CG-scale DDI complex graphs will then be leveraged by a CG-scale diffusion denoising pre-training technique developed based on the atom-scale work[6], aiming to make the CG graph encoder aware the general DDI knowledge from these samples.

Specifically, by joint modelling of protein complex conformation and underlying sequence information, the more comprehensive learning of CG DDI complex structures could be achieved. Based on this, the diffusion mechanism[10] is used to naturally add noise with varying magnitudes into the 3D coordinates and sequences of bead nodes within CG-scale DDI complex graphs to corrupt these graphs, and then to force the CG protein encoder to recover the original conformation and sequence for the complete joint modelling. The whole process is formulated as follows, the equations (6) and (7) represent the noising adding and denoising processes respectively.

$$q(D^{1:T}|D^0) = \prod_{t=1}^{T} q(D^t|D^{t-1}) \tag{6}$$

$$p_\theta(D^{0:T-1}|D^T) = \prod_{t=1}^{T} p_\theta(D^{t-1}|D^t) \tag{7}$$

in which the both processes can be decomposed into multiple time steps of noising adding or denoising following the rule of Markov chains (i.e., the total time steps are $T$). For each step $t$, $q(D^t|D^{t-1})$ and $p_\theta(D^{t-1}|D^t)$ define the current noising adding and denoising based on the parameterized CG graph encoder, respectively (in which $D^t$ represents the status of the noised CG DDI complex (graph) at step $t$). Additionally, $q(D^t|D^{t-1})$ and $p_\theta(D^{t-1}|D^t)$ can be further decomposed into the CG conformation and bead sequence noising adding and denoising processes, to realize the aforementioned joint modelling (equations (8)-(9)).

$$q(D^t|D^{t-1}) = q(S^t|S^{t-1}) \cdot q(C^t|C^{t-1}) \tag{8}$$

$$p_\theta(D^{t-1}|D^t) = p_\theta(S^{t-1}|D^t) \cdot p_\theta(C^{t-1}|D^t) \tag{9}$$

in which $C^t$ and $S^t$ denote the status the of CG DDI complex conformation and bead sequence at step $t$, respectively.

For $q(S^t|S^{t-1})$ that is performed prior to the $q(C^t|C^{t-1})$ in each step of noising adding, all side chain bead nodes $S$ (and their corresponding edges and features), which belong to the specified amino acids (AAs) to be masked, will be cropped out from the current CG DDI complex graph (to get a new cropped graph). This is for masking the sequence information that can indicate the AA types based on the pre-defined AA masking ratio value $\rho^t$. After that, the independent coordinate noise drawn based on the Gaussian distribution (controlled by the pre-defined distribution variance value $\beta^t$) will be injected into the 3D coordinate of every bead node within current graph after the above cropping, to corrupt the overall CG-scale conformation for finishing current noising adding step.

With regards to the corresponding denoising process $p_\theta(D^{t-1}|D^t)$ for CG-scale conformation and sequence recovery, the cropped graph produced based on $q(D^t|D^{t-1})$ is sent to the CG graph encoder to generate the node-level geometric representations for current corrupted CG DDI complex. These representations will be further sent to the structural denoising network and sequential denoising network, for predicting 1) the CG-scale conformation change (brought by CG conformation noise adding) measured by the distance between pairwise bead nodes of edges within the cropped graph, and 2) the AA types which are masked in CG sequence noising adding, respectively. The mean square error (MSE, denoted as $L^C$) and the cross-entropy (CE, denoted as $L^S$) are chosen as the loss function to measure the respective differences between the recovered values and corresponding ground truths. The total loss to guide the pre-training model optimization can thus be formulated as follows, in which $\alpha$ is the loss weight to balance the both denoising process.

$$L^{pre-training} = \alpha L^S + (1-\alpha)L^C \quad (10)$$

For the aforementioned AA masking ratio $\rho^t$, distribution variance $\beta^t$, structural denoising network, and sequential denoising network, we follow the settings in ref. [6], we suggest referring the original paper for the detailed description. On top of this, the pre-training is performed based on the CG graph encoder on the curated CG-scale 3DID pre-training set under 200 epochs with $\alpha = 0.5$, followed by additional 50 epochs with $\alpha = 0.8$. In addition, another difference compared with the atom-scale work in ref. [6] is that, we do not incorporate the similar conformer generation mechanism, which aims to bring more conformation varieties of original proteins, into the CG-based diffusion training process, for which we find that it does not help to the further performance improvement for our CG-scale geometric learning framework.

### Summary of implementation tools for MCGLPPI

Our basic program language is Python 3.9.18, on which Pytorch 1.12.1[11] and Torchdrug[12] 0.2.1 with a default random seed 0 is used to construct the overall framework of the MCGLPPI. For the CG-scale force field parameter generation, the Python-based MARTINI script (version 2.2, MARTINI22) martinize.py is used for the transformation process. Besides, before the parameter generation, we adopt the pdbfixer tool (https://github.com/openmm/pdbfixer) to complete missing side-chains and convert non-natural amino acids to their natural counterparts.
